# Testing vivo-morpholino mediated gene knockdown in threespine stickleback

**DOI:** 10.64898/2026.02.24.707669

**Authors:** Stella M. DiPippo, Andrea Roth-Monzón, Daniel I. Bolnick, Arshad Padhiar

## Abstract

Antisense vivo-morpholino oligonucleotides (vivo-MOs) allow transient gene knockdown in adult organisms with high specificity and low toxicity. Vivo-MOs are used in cell culture and in many established model organisms, but a method for their use has not been described in threepsine stickleback (*Gasterosteus aculeatus* (Linnaeus, 1758)). Stickleback are an emerging model system used in evolutionary and ecological genetic studies. While genomic techniques are commonly used in stickleback research, there are few studies and tools available to assess gene function in-vivo, especially for genes that may be difficult to knock out by CRISPR (e.g., lethal knock-outs). Here, we test the use of splice-blocking vivo-MOs for gene knockdown in stickleback using intraperitoneal injection of vivo-MOs targeting three candidate genes. Gene expression was assessed in the liver, spleen, and intestine. Successful knockdown of *Spi1b* was observed in the spleen, however, we observed no other significant knockdown at either timepoint tested. Injection of a fluorescently labeled control vivo-MO confirmed delivery to each target organ, validating this approach, but delivery was variable which may explain inconsistent effects. These results indicate that vivo-MOs have potential as a tool for in-vivo gene knockdown in stickleback. Optimizing delivery methods could improve reproducibility and knockdown efficiency in future studies.

## Introduction

Morpholino oligonucleotides (MOs) are stable, antisense oligonucleotides used to transiently knockdown gene expression by blocking translation or splicing (Moulton and Jiang, 2009). MOs microinjected into early embryos enter the cytosol of daughter cells and diffuse into the nucleus to bind complementary RNA sequences (Moulton, 2007). Vivo-MOs are used in adult organisms and are useful for the knockdown of genes required for survival in early development. Unlike standard MO oligos, vivo-MOs have an attached, positively charged, octa-guanidinium dendrimer group and are thought to enter cells through hydrogen bonding and electrostatic interactions with the cell membrane (Morcos et al., 2008). Vivo-MOs have successfully been used in many model species including zebrafish (*Danio rerio* (Hamilton, 1822)) (Kim et al., 2010, Shihabeddin et al., 2024), mummichog (*Fundulus heteroclitus* (Linnaeus, 1766)) (Notch et al., 2011), and mice (*Mus musculus* (Linnaeus, 1758)) (Ferguson et al., 2013), and show a potential for use in humans (Subbotina et al., 2015). However, vivo-MOs have not been used in many emerging model species, including threespine stickleback fish (*Gasterosteus aculeatus* (Linnaeus, 1758)).

Threespine stickleback are an important model for studies of evolutionary and ecological genetics (Reid et al., 2021). Marine stickleback colonized freshwater habitats in the northern hemisphere following the last glacial maximum, resulting in distinct morphological and phenotypic differences between freshwater populations and their marine ancestors (Bell and Foster, 1994). Freshwater populations also differ from one another due to adaptation to diverse ecological environments. Genomic methods such as sequencing and quantitative trait loci (QTL) mapping are commonly used to evaluate population differences and understand the genetic basis of adaptive traits in stickleback. However, while some studies have used techniques such as CRISPR and TALENs for gene knockout, in-vivo evidence for the function of candidate genes in stickleback is limited (but see Colosimo et al., 2005, Flanagan et al., 2025, Wucherpfenning et al., 2019, and Wucherpfenning et al., 2022).

Here, we evaluate the potential to use vivo-MOs to manipulate gene expression in stickleback. Compared to other gene knockdown methods such as siRNA, morpholinos have fewer off-target effects and a higher binding affinity (Summerton, 2007, Pandey et al., 2014). They are ∼25 bases in length and designed to bind complementary RNA sequences to sterically block splicing or the initiation of translation. If translation-blocking MOs are used, knockdown of protein product can be validated using a western blot (Moulton and Jiang, 2009). For splice-blocking MOs, RT-PCR is used. Depending on the location of the targeted splice junction within the coding region, splice-blocking MOs can cause exon skipping, intron inclusion, or the activation of a cryptic splice site (Moulton and Jiang, 2009). The resulting band size compared to the control splicing pattern indicates which splicing event occurred. Splice-blocking MOs can also reduce target mRNA levels, as exon skipping may introduce frameshift mutations that create premature stop codons, leading to nonsense-mediated decay (NMD) of the truncated transcript (Carrard and Lejeune, 2023). Using vivo-MOs in stickleback would enable candidate gene function to be easily determined in-vivo to validate the findings of genomic studies and identify novel gene functions. Compared to other gene modifying tools, splice-blocking vivo-MOs would be particularly useful in stickleback research, as a lack of commercial antibodies limits protein-level validation in other knockdown/knockout experiments.

To test vivo-MOs in stickleback, we targeted three genes suggested to play a role in the fibrotic immune response. Fibrosis is a defense mounted in response to parasitic tapeworm infection in stickleback, and the severity of fibrosis varies between populations (Weber et al., 2022). *Spi-1 proto-oncogene b* (*Spi1b*) encodes the transcription factor PU.1 which is present at promoters of pro-fibrotic genes (Wohlfahrt et al., 2019), and a higher expression of *Spi1b* was observed in a fibrotic fish population compared to a non-fibrotic population (Weber et al., 2022, Fuess et al., 2025). Similarly, *signal transducer and activator of transcription 6* (*STAT6*) was shown to be a strong target of natural selection in stickleback within a QTL associated with fibrosis-mediated tapeworm growth suppression (Weber et al., 2022). *STAT6* encodes a transcription factor that regulates the expression of cytokines, collagen, and fibronectin (Huang et al., 2023). *Hepatocyte nuclear factor 4-alpha* (*HNF4α*) is a third candidate gene for stickleback immunity to *Schistocephalus* tapeworms, also a strong target of selection within a QTL for resistance, and identified as the origination of a cascade of differential expression in stickleback exposed to *Schistocephalus solidus* (Müller, 1776) infection (Weber et al., 2022). *HNF4α* encodes a transcription factor that regulates liver gene expression and decreases fibrosis in human hepatocytes (Yeh et al., 2019). We selected *Spi1b*, *STAT6, and HNF4α* as candidate genes to test the efficiency of vivo-MO mediated gene knockdown in stickleback, which could enable future functional characterization of these, or other, candidate genes.

## Materials and methods

### Stickleback fish collection

Threespine stickleback (*G. aculeatus*) were collected from Sayward Estuary, Echo Lake, and Boot Lake on Vancouver Island, British Columbia. Fish were collected with a scientific fish collection permit from British Columbia (permit number NA23-787881). Hybrid crosses between Boot x Echo and Sayward x Echo parents were generated in June 2022 and raised to maturity. F1 progeny of the Boot x Echo cross were used in the morpholino knockdown experiments. F1 Sayward x Echo progeny were used in a fluorescent injection experiment to test MO delivery efficacy. Protocols were approved by the University of Connecticut Institutional Animal Care and Use Committee (IACUC protocol number A24-017), and field collection permits were provided by the Ministry of the Forests Lands and Natural Resources, British Columbia (NA22-679623).

### Morpholino Experiment

#### Morpholino design

We designed splice-blocking vivo-MOs for S*TAT6*, *Spi1b*, and *HNF4α* which were then synthesized by Gene Tools (Philomath, Oregon, USA) (Table 1). Each vivo-MO was designed to bind complementary sequences at internal splice junctions. The S*TAT6-* and *HNF4α*-MOs span exon-intron boundaries, while the *Spi1b*-MO binds an intron sequence directly adjacent to an internal exon. MO binding in these regions blocks splicing machinery and should cause skipping of the targeted exon, resulting in a truncated transcript. A fluorescein tagged vivo-MO from Gene Tools was used as a standard control. The lyophilized MOs were dissolved in water to create 0.5 mM stock solutions which were stored at room temperature.

**Table 1:**
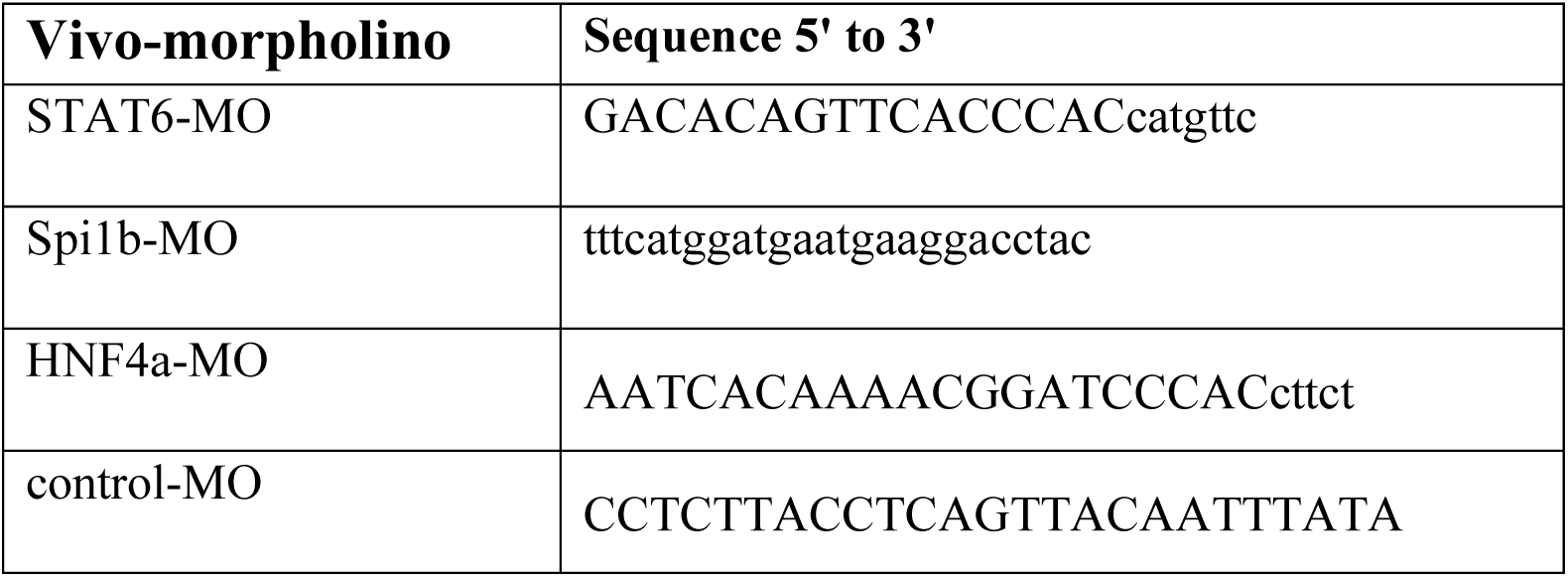
Vivo-morpholino sequences. Exon and intron sequences are denoted by capitalized and lowercase letters, respectively.

#### Morpholino injection

Adult Boot x Echo F1 hybrid stickleback were anesthetized in neutral buffered 0.14 g/L MS-222, then 15 μl of 0.125mM (in PBS) vivo-MO solution was injected intraperitoneally into each fish using an 8mm 31G insulin syringe (MHC medical products, Ohio, USA). After injection, the fish were placed in a 22 °C tank (from a 17 °C tank) to increase uptake, vivo-MOs being designed for use in higher body temperature organisms. In the first injection experiment, the vivo-MOs were each injected into two fish each and the fish were dissected 48 hours post-injection (hpi). In the second injection experiment, each vivo-MO was injected into three fish each and the fish were dissected at 24 hpi.

#### RNA isolation and semi-quantitative RT-PCR

After injection, at the specified timepoint, the fish were euthanized in MS-222 and the spleen, liver, and intestine were removed and stored in RNAlater at 4 °C. The tissue was homogenized using ceramic beads, then the RNA was isolated using the QIAGEN RNeasy PlusMini Kit, following the manufacturer’s protocol (Qiagen, Hilden, Germany). Complementary DNA (cDNA) was synthesized from 1 μg of RNA using the Bio-Rad iScript cDNA synthesis kit (Bio-Rad, Hercules, California, USA), following the manufacturer’s protocol.

PCR was used to assess gene knockdown. Promega 5X green GoTaq Reaction Buffer (Promega, Wisconsin USA), 5 μM each of forward and reverse primers (Table 2), and 2.5 μl of whole or 1:10 diluted (in water) template cDNA were used. Initial denaturation occurred at 95 °C for 2 minutes, followed by 32 cycles (28 cycles for reference genes) of 95 °C for 15 seconds, 55 °C for 15 seconds, and 72 °C for 30 seconds. A final extension occurred at 72 °C for 5 minutes.

**Table 2:**
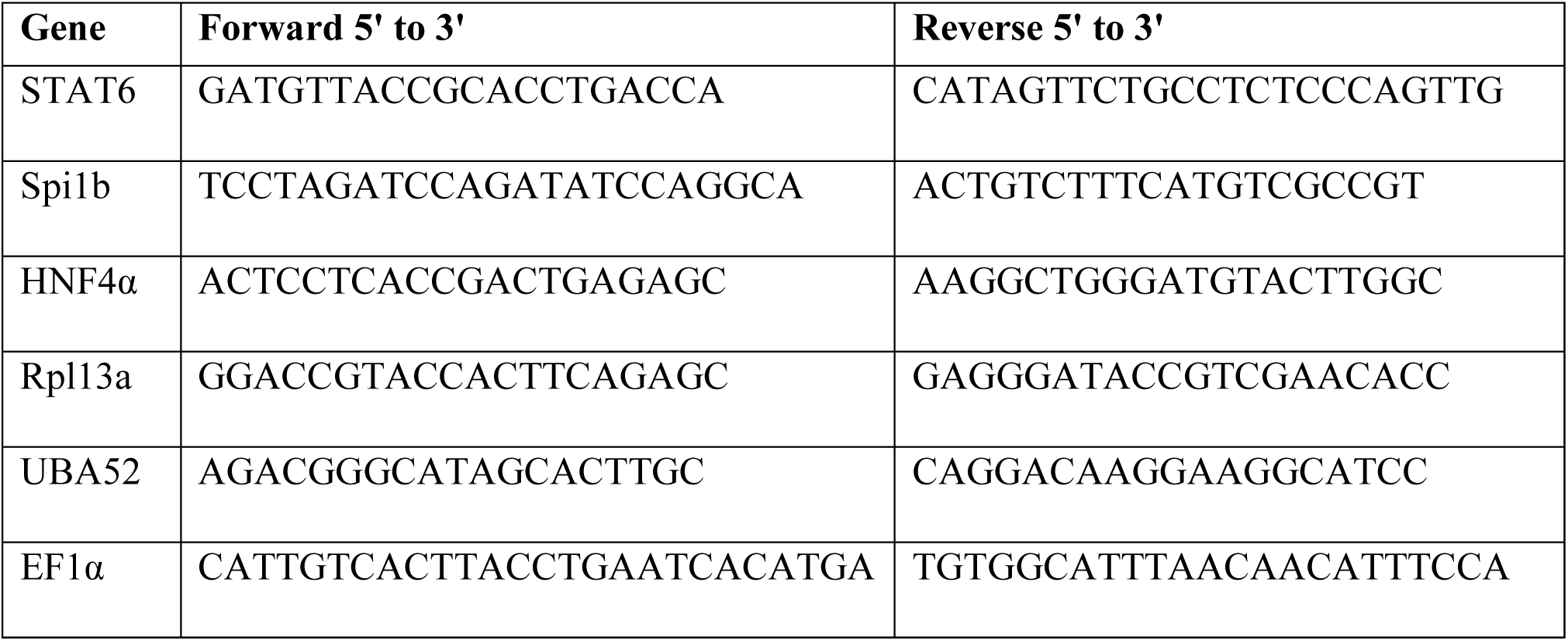
PCR primer sequences for evaluation of gene knockdown. *UBA52* and *EF1α* sequences are from Hibbeler et al., 2008, and McCairns et al., 2009, respectively.

Primers for the evaluation of gene knockdown were designed to bind exons upstream and downstream of the MO targeted splice sites. Targeting internal splice junctions causes exon skipping and should therefore result in a shorter transcript on a gel in MO injected fish compared to controls. NMD of the truncated transcript is also possible and would cause a fainter or undetectable wild-spliced band on the gel (Carrard and Lejeune, 2023). RT-PCR products were visualized on a 2.5% agarose gel and band intensities were quantified using ImageJ (ImageJ, National Institutes of Health, USA).

### Fluorescent control vivo-MO injection

To determine the efficiency of delivery and cellular uptake of the vivo-MOs at different timepoints, we intraperitoneally injected twelve fish with 15 μl of 0.1 mM (in PBS) fluorescent vivo-MO Std control (Gene Tools), following the injection protocol described above. After injection, the fish were returned to a 17 °C tank. At each timepoint (24, 48, 72, and 96 hpi), three fish were euthanized in MS-222, then the spleen, liver, and intestine were removed and placed on ice. The organs were fluorescently imaged within 30 minutes of dissection using an IVIS Spectrum (PerkinElmer; Hopkinton, MA, USA) with FITC settings (excitation around 465–495 nm and emission around 515–575 nm). A non-injected control organ was included in each imaging session, and fluorescence was quantified relative to the control. Organs were placed on a non-reflective black background to minimize autofluorescence. The IVIS Spectrum can also be used for whole-fish CT scanning (Supplementary video S1), which may enable visualization of internal parasites and other structures in live fish without the need for euthanasia.

### Statistical analysis

Statistical analysis was performed using Microsoft Excel. Standard error was calculated for each group with more than one fish. To determine the significance of gene knockdown, t-tests were used to compare mean gene expression in control and experimental vivo-MO injected fish. A p-value < .05 indicates significant gene expression differences between control and experimental groups.

## Results

### Morpholino experiment

#### Vivo-MO mediated knockdown of Spi1b in the spleen at 24 hpi

Twelve fish were injected with either a control or experimental vivo-MO and gene knockdown was assessed at 24 hpi. One *Spi1b*-MO injected fish died by 24 hpi, which could reflect either an injection injury or MO toxicity. Gene expression was analyzed in the liver and spleen of surviving individuals. mRNA transcripts in the spleen appeared at the same band size in both control and morphant fish (Figs. 1A and B), indicating either that any truncated transcripts were degraded by NMD, or that no exon skipping occurred. Exon skipping without NMD would have resulted in shorter bands present on the gel. Expression of *Spi1b* in the spleen decreased by greater than 50% (t = 43.2, p < 0.0001) in *Spi1b-*MO injected fish compared to controls (Fig. 1D), indicating successful knockdown. *Spi1b* knockdown was observed as a reduced intensity of the band at the expected WT splice pattern size (Fig. 1B). *STAT6* expression increased 1.1-fold (t = -3.25, p = .042) in the spleen of *STAT6*-MO injected fish compared to controls (Fig. 1C), which was unexpected, but could reflect a compensatory response. No significant changes in expression of *STAT6* (t = -4.95, p = 0.99), *Spi1b* (t = -0.56, p = 0.69), or *HNF4α* (t = 0.74, p = 0.25) were observed in the liver between MO and control fish at 24 hpi (all p > 0.05; data not presented graphically). The Excel file used to calculate statistics and summary results is included as Supplementary (Table S1).

**Figure 1:**
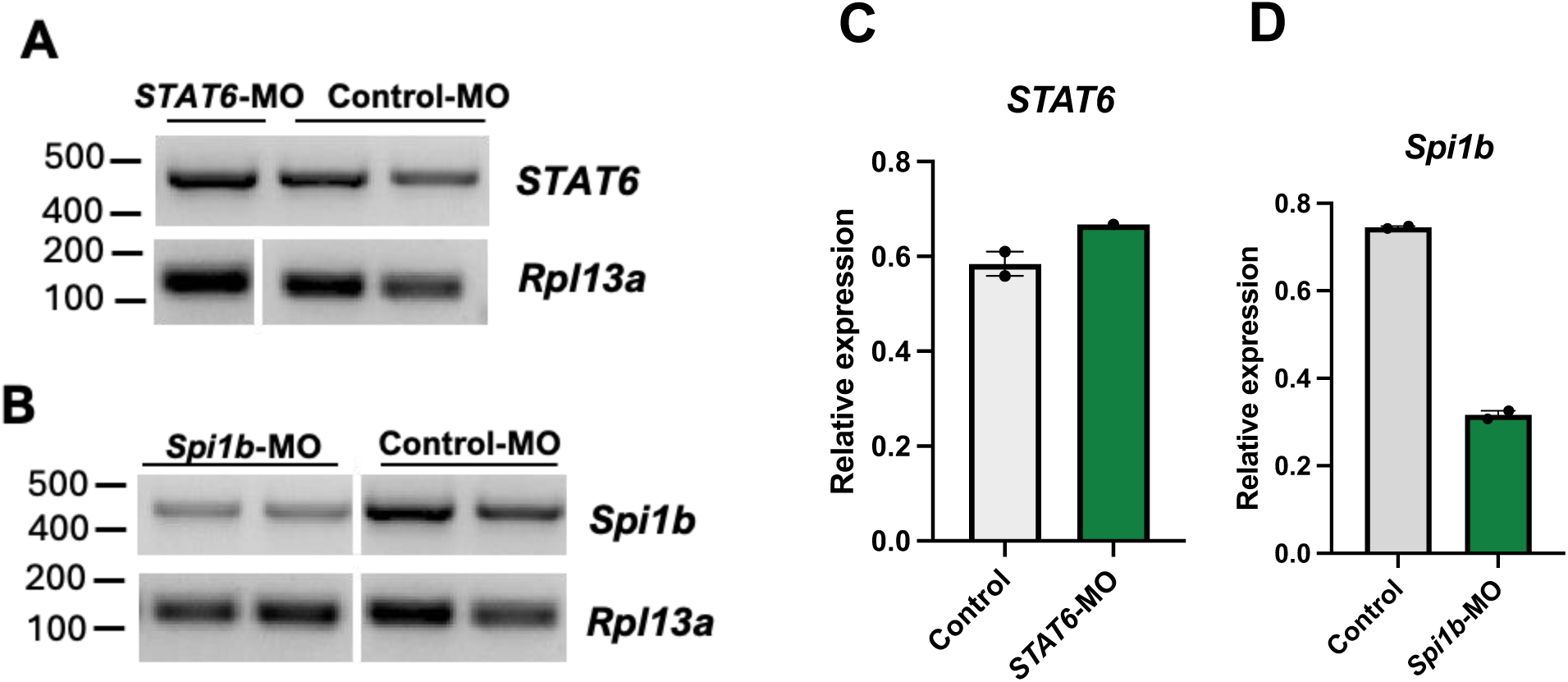
Knockdown of *Spi1b* expression in the spleen at 24 hpi. (A and B) Semi-quantitative RT-PCR analysis of gene expression in control, (A) *STAT6-*MO, and (B) *Spi1b-*MO injected fish. Non-adjacent lanes from the same gel were rearranged as indicated by vertical white lines. (C and D) Band intensities were quantified using ImageJ and normalized to *Rpl13a*. Values are mean ± SE. The absence of error bars for the *STAT6*-MO data is because N=1.

#### Morpholino mediated gene knockdown in the intestine at 48 hpi

To investigate the efficiency of vivo-MO mediated knockdown at a later timepoint (48 hpi), an additional eight fish were injected with a control or experimental vivo-MO. One *STAT6-*MO and one *HNF4α*-MO injected fish died by 48 hpi. Gene expression was analyzed in the intestine and liver of surviving individuals. As seen by the absence of target gene bands in morphant fish (Figs. 2A and C), complete knockdown of *STAT6* and *HNF4α* was observed in the intestine when expression was normalized to *EF1α* (Eukaryotic translation elongation factor 1 alpha) (Figs. 2D and F), though this was amendable to statistical confirmation because the mortalities led to N=1 sample sizes per treatment. The reference genes *Rpl13a* (Ribosomal protein L13A) and *UBA52* were also used, however, little to no expression of either reference gene was detected in three of the four morphant fish (Figs. 2A-C), suggesting potential off-target effects of the MOs. There was no significant change in *Spi1b* expression in the intestine (Fig. 2E, t = -0.68, p = 0.718). *STAT6*, *HNF4α,* and *Spi1b* expression in the liver were not significantly different between morphant and control fish (all p > 0.05 or N = 1; data not presented graphically).

**Figure 2.**
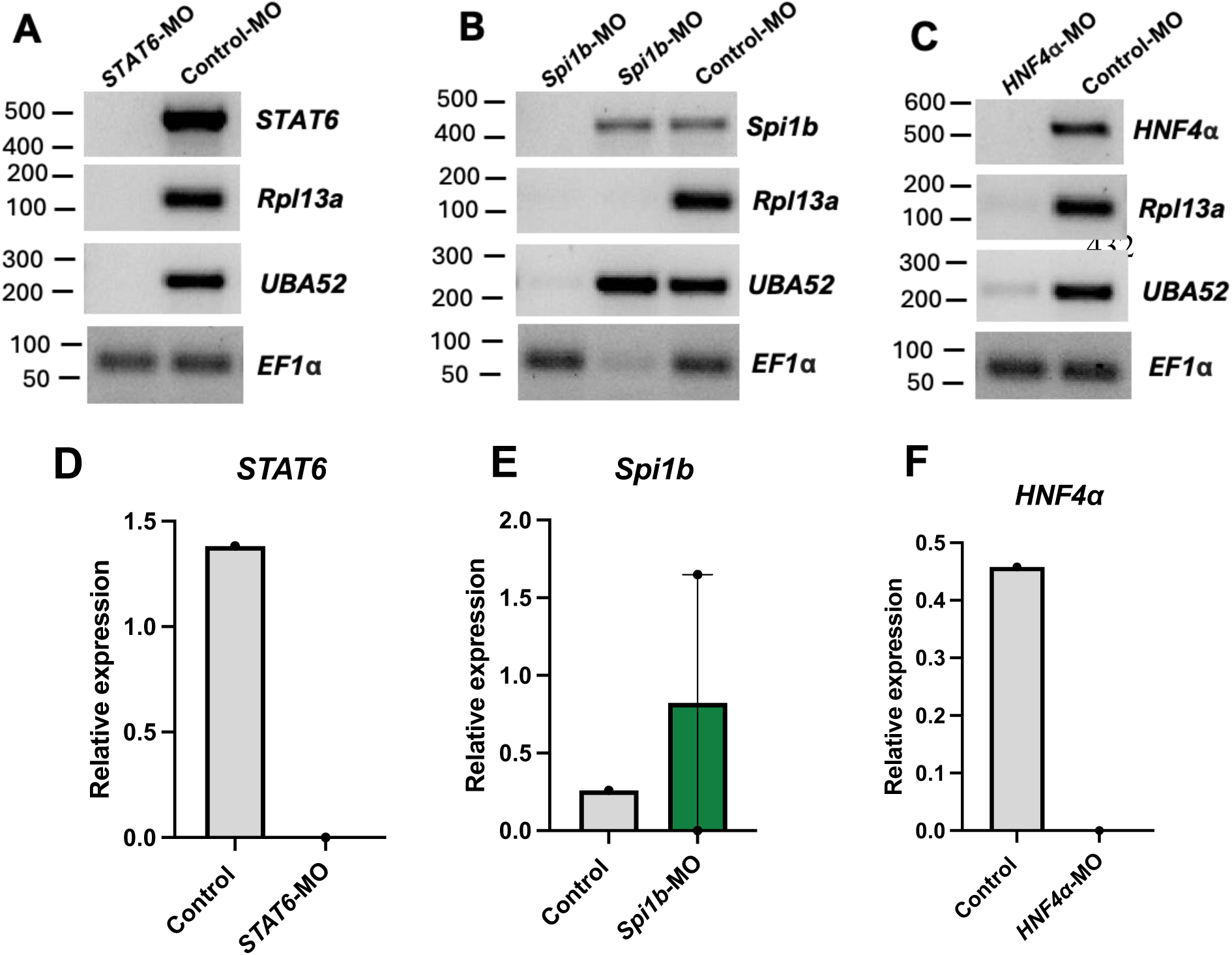
Knockdown of *STAT6*, *Spi1b*, and *HNF4α* expression in the intestine at 48 hpi. (A-C) Semi-quantitative RT-PCR analysis of gene expression in control and (A) *STAT6-*MO, (B) *Spi1b-*MO, and (C) *HNF4α*-MO injected fish. (D-F) Band intensities were quantified using ImageJ and normalized to *EF1α*. Values are mean ± SE. The lack of error bars for all but *Spi1b*-MO is because N=1.

### Fluorescent control vivo-MO injection

To determine the optimal timepoint for vivo-MO uptake, we injected 12 fish with a fluoresceinated control MO and compared the fluorescent intensity at 24, 48, 72, and 96 hpi. In the liver, fluorescence was detected at each timepoint but was strongest at 24 hpi (Fig. 3). In the spleen, the strongest fluorescence was seen in one sample at 96 hpi (Fig. 4). Minimal fluorescence was detected at 72 hpi, and there was no fluorescence observed at 48 hpi (Fig. 4). Fluorescence was detected at every timepoint in the intestine (Fig. 5). Overall, these results demonstrate inconsistent MO absorption and retention across organs and timepoints following intraperitoneal injection, which likely contributes to the variable knockdown efficiency observed in gene expression analyses.

**Figure 3.**
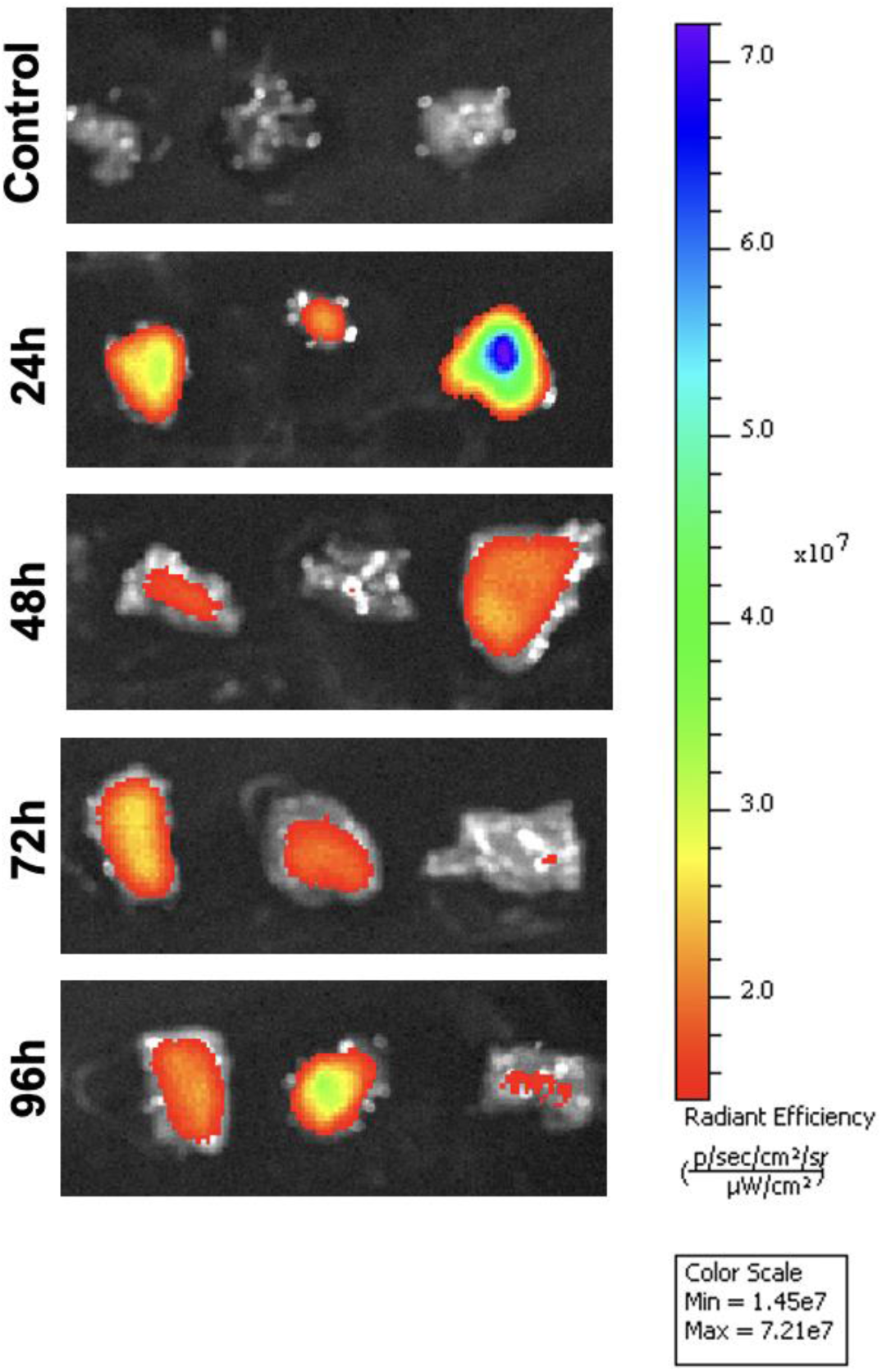
Vivo-morpholino uptake in the liver. In-vivo fluorescent imaging of a fluorescein tagged control morpholino in the liver at 24, 48, 72, and 96 hours post-injection (hpi). Livers from three individual fish are shown at each timepoint. Fluorescence intensity varied between samples, with notably low or absent uptake in replicates at 48, 72, and 96 hpi.

**Figure 4.**
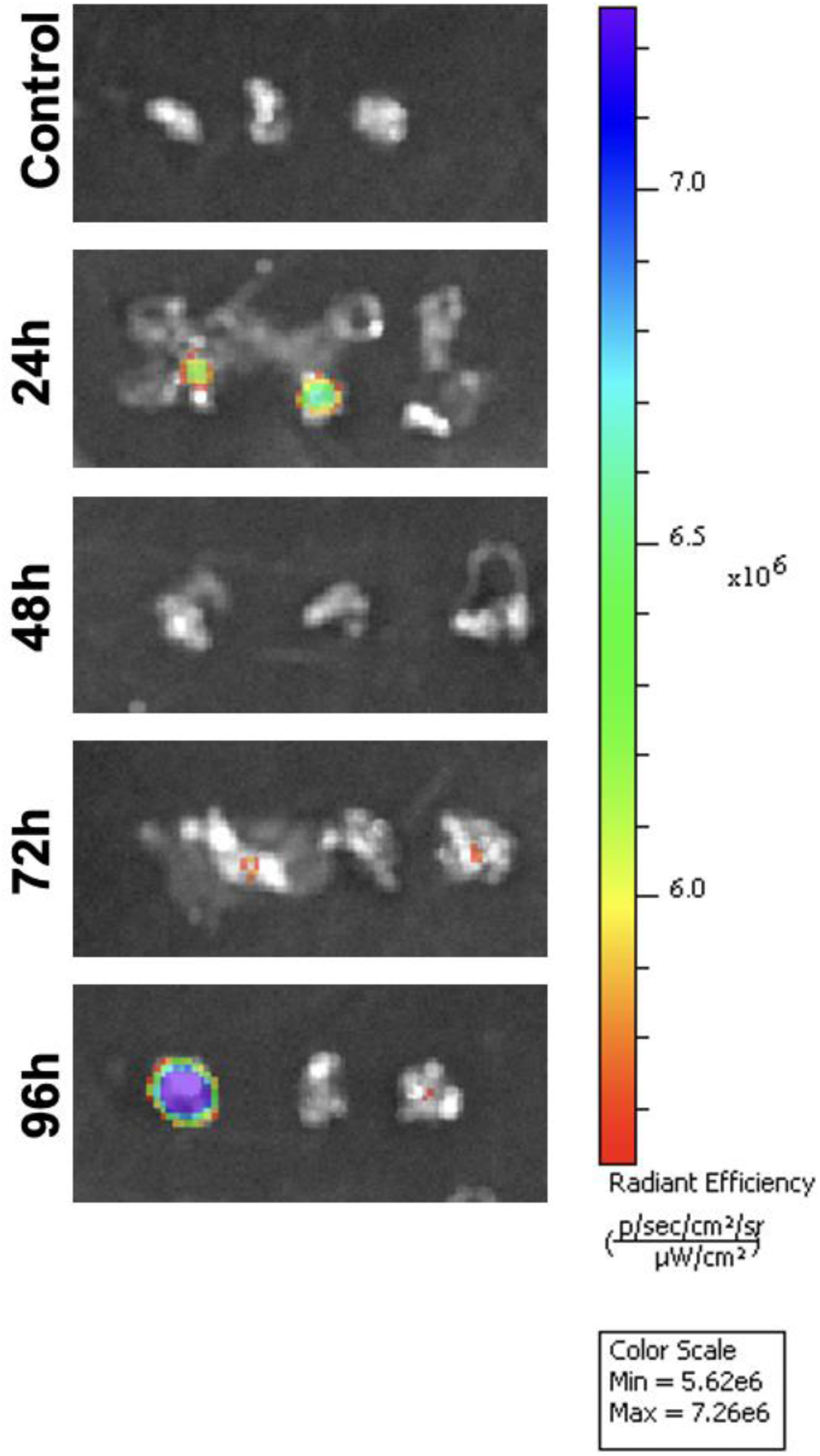
Minimal vivo-morpholino uptake in the spleen. In-vivo fluorescent imaging of a fluorescein tagged control morpholino in the spleen at 24, 48, 72, and 96 hours post-injection. Spleens from three individual fish are shown at each timepoint.

**Figure 5.**
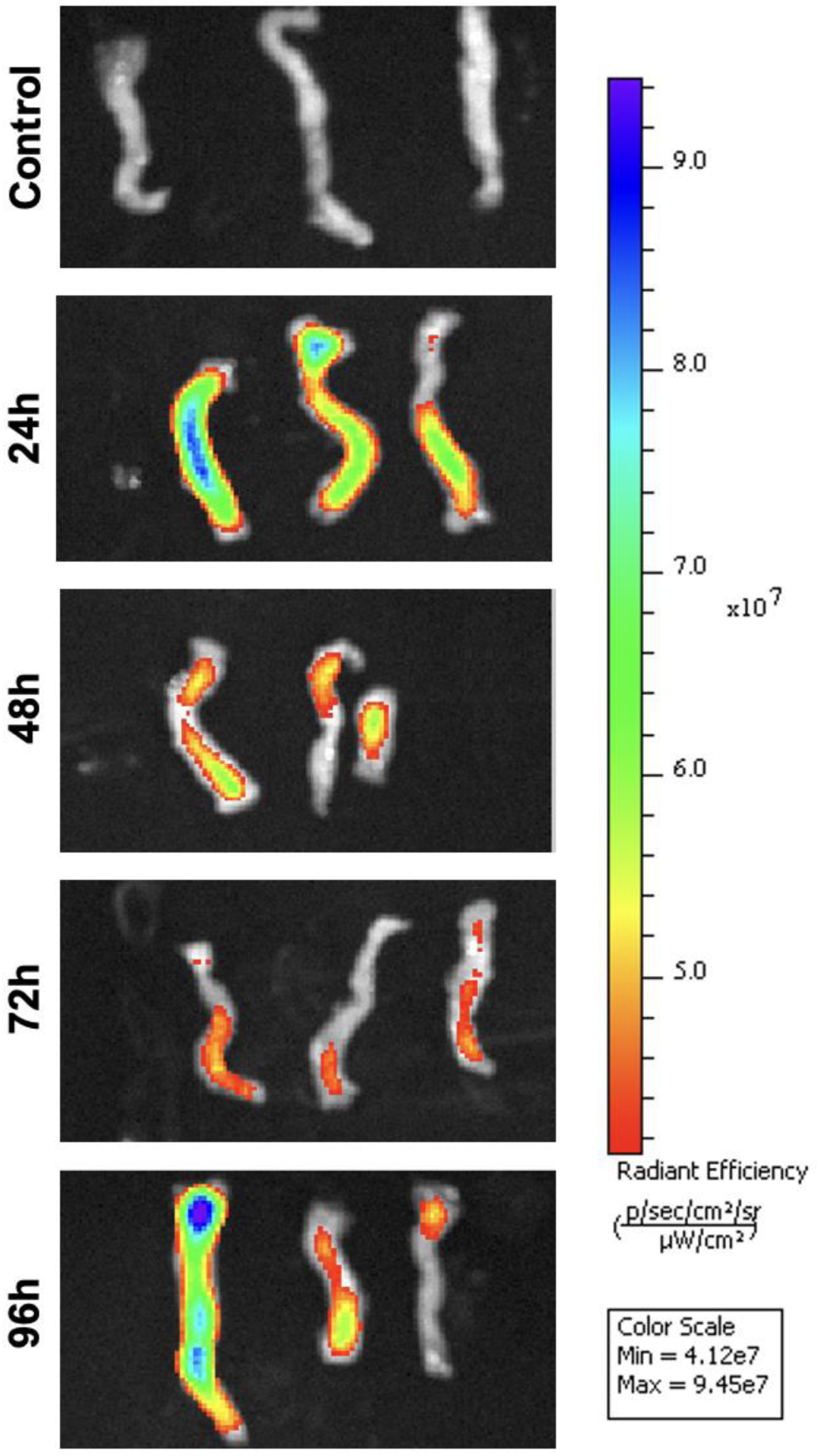
Vivo-morpholino uptake in the intestine. In-vivo fluorescent imaging of a fluorescein tagged control morpholino in the intestine at 24, 48, 72, and 96 hours post-injection. Intestines from three individual fish are shown at each timepoint. Fluorescence was detected at all timepoints, consistent with higher intestinal knockdown efficiency, but showed marked variability between replicates, indicating inconsistent delivery and retention despite direct peritoneal exposure.

## Discussion

Vivo-MOs are a useful tool for gene knockdown in many model species (Kim et al., 2010, Shihabeddin et al., 2024, Ferguson et al., 2013, Notch et al., 2011), but there is currently no protocol for their use in threespine stickleback. Stickleback are an emerging model system important for evolutionary genomic research. However, studies that evaluate phenotypic gene information in stickleback are limited, typically requiring gene editing in embryos that can be difficult for highly pleiotropic or essential genes, such as *Spi1b* (Flanagan et al., 2025). Vivo-MOs would provide a cost-effective and simple method to transiently knockdown genes in adult stickleback in a targeted tissue. We tested vivo-MOs targeting three candidate genes suggested to be involved in the fibrotic immune response in stickleback. We examined knockdown in the spleen, intestine, and liver following intraperitoneal injection of vivo-MOs. Future work will build on these results to test these knockdown effects on the fibrosis response to infection or antigen injection.

Morpholino mediated gene knockdown is transient and decreases overtime, as seen in *F. heteroclitus* (Notch et al., 2011). We evaluated gene knockdown at 24 hpi and saw a significant decrease in *Spi1b* expression in *Spi1b*-MO injected fish compared to controls (Fig. 1D). The mRNA transcripts in both the control and *Spi1b*-MO injected fish were equal size, but the morphant fish had quantitatively dimmer *Spi1b* bands, suggesting that NMD of any truncated transcripts occurred. We did not observe significant knockdown of any target gene in the liver at 24 hpi. This may be due to insufficient volumes of the vivo-MO solution reaching the liver, decreased cellular uptake of the vivo-MOs, or its rapid degradation in the liver.

A limitation of using MOs, as with other gene modifying techniques, is the potential for downstream effects or non-specific binding that affects the expression of non-target genes. We observed the complete absence of target gene expression in the intestine at 48 hpi, however, these samples had little to no expression of two of the tested housekeeping genes (but not all the housekeeping genes, indicating that overall RNA was of sufficient quality). Though this result could be due to partial RNA degradation. Similar variability across individuals and genes suggests uneven morpholino distribution rather than uniform knockdown efficiency. The apparent loss of signal in the intestine may reflect localized exposure following intraperitoneal injection rather than consistent gene-specific suppression. This proximity may also contribute to variability and the observed effects on reference genes. In future studies, housekeeping genes (e.g., those with consistent expression across cell types and environmental conditions) which are unaffected by the MOs should be used to evaluate knockdown by splice-blocking MOs.

Of each gene, timepoint, and organ tested, we only observed significant knockdown of *Spi1b* in the spleen at 24 hpi. This inconsistent knockdown pattern likely stems from IP delivery limitations (Dedrick and Flessner, 1997). For effective knockdown, vivo-MOs must avoid leakage back out through the injection wound (a problem with stickleback skin), reach target organs, enter cells, then bind complementary RNA. If not delivered in a sufficient amount, too little vivo-MO may reach individual organs and result in little to no uptake.

As a proof-of-concept experiment to ensure delivery of vivo-MOs at the timepoints tested, we injected a fluoresceinated control vivo-MO and observed fluorescence in each organ at 24, 48, 72, and 96 hpi. Fluorescent intensity was often strongest at earlier compared to later timepoints (Figs. 3 and 5), suggesting the amount of MO in cells decreased over time; however, there was variation between samples. This could be due to slight differences in injection volume and location in each fish, or varying leakage at the injection wound. Intra-venous (Matrone et al., 2021) and tissue specific injection (Shihabeddin et al., 2024, Hyde et al., 2012) are additional methods of vivo-MO delivery used in other models that may improve consistency and delivery to targets organs in future studies.

We sought to develop a method for in-vivo gene knockdown in stickleback using splice-blocking vivo-MOs to target *STAT6*, *Spi1b*, and *HNF4α.* We observed significant knockdown of one gene in the spleen, while intestinal signal changes were variable and difficult to interpret confidently. Future studies should test increasing concentrations of MO while minimizing lethality, and alternative delivery methods in stickleback. Vivo-MOs could provide an effective tool for gene knockdown in adult stickleback to evaluate the findings of genomic studies and assess gene function.

## Acknowledgments

We thank the University of Connecticut In Vivo Imaging System Core facility for use of their IVIS equipment. We thank Vince Vacco for animal care, and Lauren Simonse and Lindsay Yue for assistance with field work to collect adult fish and generate the crosses for this research.

## Author contribution statement

The research presented here was conceived of by AP in consultation with DIB and SMD. The data collection was conducted by SMD and AP. Animal care and rearing of the fish was done by ARM. Analyses were done by SMD, and writing was completed by SMD with feedback from DIB, ARM, and AP.

## Data availability statement

Data is available in article supporting information.

