## Supplementary Video S1 for "Testing vivo-morpholino mediated gene knockdown in threespine stickleback"

### Slide 1
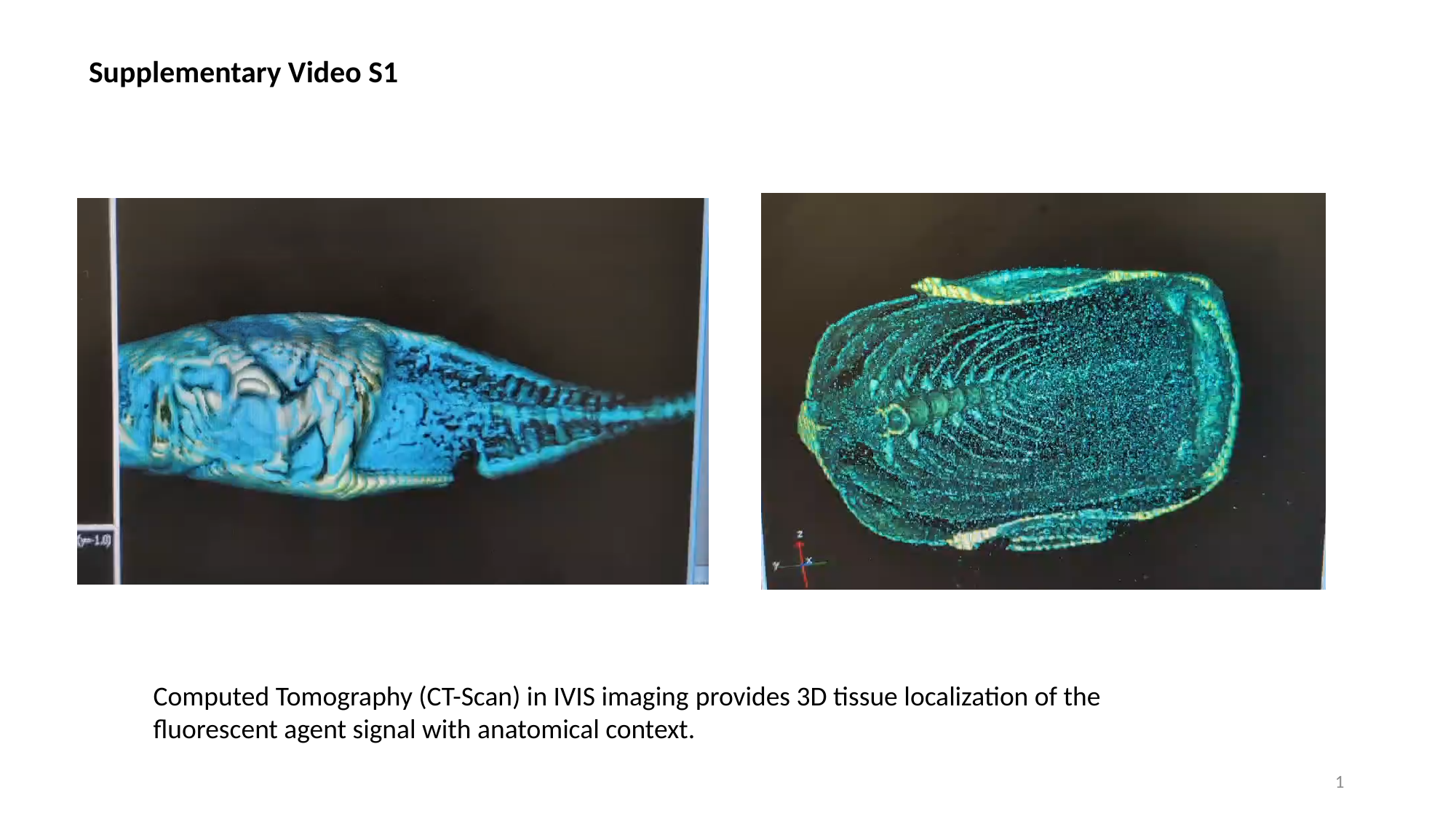

Supplementary Video S1
Computed Tomography (CT-Scan) in IVIS imaging provides 3D tissue localization of the fluorescent agent signal with anatomical context.
1
